# Unraveling the Phenotype of Dormant Metastases Controlled by the Immune System

**DOI:** 10.1101/2024.12.02.626440

**Authors:** Virginia Chamorro, Ignacio Algarra, Verónica Sanz, María Pulido, Irene Romero, Estefanía Chico, Marina Millán, María Escaño-Maestre, Isabel Linares, Ángel M. García-Lora

**Affiliations:** Servicio de Análisis Clínicos e Inmunología, UGC Laboratorio Clínico, Hospital Universitario Virgen de las Nieves, Granada, Spain; Instituto de Investigación Biosanitaria ibs.GRANADA, Granada, Spain; Departamento de Ciencias de la Salud, Universidad de Jaén, Jaén, Spain; Servicio de Análisis Clínicos, Hospital de Antequera, Málaga, Spain; Plataforma Biobanco ibs.Granada, Nodo Granada, Granada, Spain

**Keywords:** Dormant metastases, Antitumor immunity, gene expression, chemokines, Biomarkers, immune microenvironment

## Abstract

During the progression of cancer, metastatic cells frequently enter a dormant phase. The immune system plays a crucial role by restricting the growth of dormant metastases, although it cannot eradicate them. In our laboratory, we developed a preclinical mouse model of metastatic dormancy. Dormant spontaneous metastases are controlled by the immune system of wild-type mice. Depletion of the host immune system causes these metastases to awaken and progress. The elucidation of the phenotype, genes, miRNAs and immune cells involved in the phenomenon of metastatic dormancy may prove in the development of effective strategies for combating metastatic disease. Dormant Metastases are compared with Nude Metastases and Overt Metastases that have never been in dormancy. The findings of the study indicate that the dormant metastases exhibit a unique and differentiated phenotype. This is evidenced by their varied response to nutrient-restrictive conditions, chemotherapeutic agents, and cytokines *in vitro*. Additionally, dormant metastases display a unique pattern of gene and miRNA expression. The microenvironment of dormant metastases shows an increase in certain immune cell subpopulations. Immune-controlled dormant metastases exhibit a unique phenotype that can be exploited to discover new biomarkers, as well as to develop therapies to eradicate them or control overt metastases.

## 1 INTRODUCTION

The main challenge in fighting cancer is destroying metastases, which are the primary cause of death in most cases (1). The progression from primary tumor to metastasis involves several stages, including tumor vascularization, intravasation into blood vessels, extravasation and dissemination to other organs (2, 3). However, when tumor cells reach new organs, they can enter a period of latency known as dormant metastasis (4). Metastatic cells remain dormant during this period, with minimal or no proliferation, and are not destroyed (5, 6). This period can last for years or even decades after the primary tumor has been removed, as is often observed in cases of breast and prostate cancer (7, 8). Metastatic dormancy can occur due to anti-angiogenic factors (angiogenic dormancy), proliferation-suppressing factors (cellular dormancy), or the action of the immune system (immune dormancy) (9, 10). These three types of latency are interrelated and may all play a role in the process of metastatic dormancy (11).

During immune dormancy, metastatic growth is suppressed by the immune system, but the immune system is unable to eliminate dormant metastatic cells (12, 13). Further research could investigate the phenotype of dormant metastases, as well as the molecular mechanisms and immune cells involved in maintaining this equilibrium (14). By understanding the immune mechanisms involved in this balancing process, we may be able to manipulate the immune system to destroy dormant metastases before they become overt. We may also be able to control overt metastases in a state of immunodormancy (13).

Additionally, understanding the phenotype of dormant metastases may aid in the detection and discovery of new biomarkers of dormant metastatic disease (15).

Studying dormant metastases in humans is a challenging task since they are undetected and cannot be isolated for study. Therefore, preclinical murine models are necessary to accurately replicate the process of metastatic dormancy (16, 17). Furthermore, it is essential to develop models that accurately depict every stage of tumor progression toward metastatic dormancy. The use of spontaneous metastasis assays in murine preclinical models is the only way to fully replicate these stages. Internationally, there are very few preclinical murine models with the described characteristics. Our laboratory has developed a preclinical murine model of spontaneous immune-mediated dormant metastasis in wild-type mice without the use of any additional treatments (18). Methylcholanthrene was used to induce a primary tumor in BALB/c mice. The tumor was then removed, cultured, and cloned by limiting dilution, resulting in different cell lines from these tumor clones (19). One of the clones was subcutaneously injected into BALB/c mice. Following the removal of the primary tumor, the mice were monitored for the emergence of spontaneous metastases. The study revealed that the mice did not develop overt spontaneous metastases (18). However, in immunodeficient BALB/c nude mice, all the mice developed overt spontaneous metastases. In a new assay, this tumor clone was injected into immunocompetent wild-type mice. The primary tumor was removed, and the mice were observed for 5 months. During this time, no overt metastases were detected. Next, the hosts were depleted of CD4+ and CD8+ T lymphocytes for a period of three months. Finally, the presence of overt spontaneous metastases was revealed through euthanasia of the animals. These metastases, called Dormant-met, had been in a state of immune metastatic dormancy for more than five months and were not awakened until the depletion of the host’s T lymphocytes. These metastases should exhibit a phenotype comparable to that of dormant metastases and were cultured.

In this study, we selected two of these dormant metastases, Dorman-met, and compared their phenotype, gene and miRNA expression, and different biological characteristics with two metastases generated in nude mice, Nude-met (which were not in a state of immune dormancy) (18), and with two other overt spontaneous metastases, Overt-met, produced from another tumor clone of the same primary tumor (20) (which also were not in a state of immune dormancy). The results show that these dormant metastases have a distinct phenotype in comparison to Nude-met and Overt-met. Additionally, the immune populations involved in the process of metastatic immune dormancy were analyzed.

## 2 MATERIALS AND METHODS

### 2.1 Animals

Eight-week-old male or female BALB/c mice (Charles River Laboratories) were used in the experiments. This study was carried out in accordance with the recommendations of the European Community Directive 2010/63/EU and Spanish law (Real Decreto 53/2013) on the use of laboratory animals, and their housing and the experimental procedures were approved by the Junta de Andalucía animal care committee and adhered to the animal welfare guidelines of the National Committee for Animal Experiments. The animals were anesthetized with 0.04 mL of diazepam (Valium, Roche) and 0.1 mL of ketamine (Ketolar, Pfizer). When clear signs of disease were observed, the animals were anesthetized and euthanized by cervical dislocation, followed by complete necropsy.

### 2.2 Cell lines and culture conditions

The GR9 cell line originates from a mouse fibrosarcoma induced by methylcholanthrene in BALB/c mice and has been extensively characterized in our laboratory. It is composed of cell clones with distinct H-2 class I expression patterns and metastatic capacities (19). Two specific clones, GR9-B11 and GR9-A7, were isolated from the GR9 cell line using a limited dilution method. The GR9-B11 and GR9-A7 cell lines were recloned by picking up individual cells under phase-contrast microscopy. Spontaneous metastasis assays were performed with these two GR9 cell clones (18, 20). The Dormant-met 1 and Dormant-met 2 cell lines were derived from spontaneous lung metastases induced by the GR9-B11 clone in immunodepleted wild-type BALB/c mice (18). Similarly, the Nude-met 1 and Nude-met 2 cell lines were obtained from spontaneous lung metastases induced by the GR9-B11 clone in nude BALB/c mice (18). Overt-met 1 and Overt-met 2 were derived from spontaneous lung metastases induced by the GR9-A7 clone in wild-type BALB/c mice (20). All cell lines were characterized by PCR assays using short tandem repeat analysis and regular testing for MHC-I genotype and surface expression. The cell lines were tested for Mycoplasma by PCR and were negative as of January 2024. All cell lines were used for less than 10 passages after thawing. The cell lines were cultured in Dulbecco’s modified Eagle’s medium (Sigma−Aldrich), supplemented with 10% FBS (Life Technologies), 2 mmol/L glutamine (Sigma−Aldrich), and antibiotics. In some assays, cells were cultured with 0.5% FBS for 5 days.

### 2.3 Cell seeding and proliferation assessment

A total of 3 × 10^5^ or 1 × 10^6^ tumor cells were seeded in T-25 or T-75 cell culture flasks. Daily cell counts were performed over 1–4 days using the Trypan blue exclusion method with a hemocytometer. Two independent investigators conducted the cell counts. The cell proliferation rate was determined as the ratio of the final cell number to the initial cell number.

### 2.4 *In vitro* treatment with chemotherapeutic agents and cytokines

For *in vitro* treatment assays involving chemotherapeutic agents, a dose−response curve was established. Concentrations of each chemotherapeutic were selected to achieve 100% inhibition of proliferation in the Overt-met group. The chosen concentrations for 72 hours of treatment were as follows: 200 nM cisplatin; 10 nM doxorubicin; 10 nM metotrexate; and 30 nM paclitaxel(Sigma−Aldrich). In specific experiments, cell lines were subjected to *in vitro* treatment with 50 U/mL IFN-ɣ, 12 pg/ml TNF-⍺, or 1 ng/ml TGF-β for a duration of 48 hours (Miltenyi Biotech).

### 2.5 RNA sequencing (RNA-Seq)

Total RNA was extracted using PureLink RNA Mini Kits (Invitrogen), and any remaining DNA was removed using a DNA-free DNA removal kit (Invitrogen). The quality and concentration of RNA were assessed using a QuBit 4 Fluorometer and associated kits (Thermo Fisher Scientific) and an Agilent 2100 Bioanalyzer (Agilent Technologies). The preparation of the RNA library and subsequent transcriptome sequencing were conducted by Novogene Co., Ltd. (Beijing, China). Differential expression criteria: Genes with an adjusted p-value < 0.05 and a |log2(fold change)| > 1 were considered differentially expressed. Service Provider: Novogene performed the read alignment, quality control, differential expression analysis, and pathway analysis using a standard pipeline. RNA-Seq data analysis provides valuable insights into the transcriptomic landscape, allowing the identification of differentially expressed genes and potential pathways involved in the experimental conditions studied.

### 2.6 RT−PCR and quantitative real-time PCR

For RT−PCR, 1 μg of total RNA was converted to complementary DNA (cDNA) using the iScript gDNA clear cDNA Synthesis kit (Bio−Rad) in a 20 μl reaction volume. The resulting cDNAs were diluted to a final volume of 100 μL. The real-time PCR assay was performed in a CFXConnect Real-time PCR detection system (Bio-Rad). One microliter of diluted cDNA was used per reaction. PCRs were conducted in quadruplicate, and the values obtained are expressed as the means ± SDs. Genes with an adjusted p-value < 0.05 and a fold change > 2 were considered to be differentially expressed. Quantitative PCR was conducted with the SsoAdvanced Universal SYBR Green Supermix (Bio-Rad). The PCR conditions included 40 cycles of 15 s of denaturation at 95°C and 30 s at 60°C. The following predesigned PCRPrime arrays (Bio-Rad) were utilized: Lipoprotein Signaling and Cholesterol Metabolism (SAB Target List) M96 (10034358); Chemokines and Receptors (SAB Target List) M96 (10034295); and Cell Surface Markers (SAB Target List) M96 (10034292).

### 2.7 Determination of cytokine/chemokine levels

A Bio-Plex Pro™ Mouse Chemokine Panel 33-Plex, a Bio-Plex Pro™ Mouse MCP-2 single-plex, and Bio-Plex Pro™ Mouse TGF-β 3-Plex assays (Bio-Rad) were utilized to profile the concentrations of mouse cytokines and chemokines in the cell culture supernatants. The assays were conducted following the manufacturer’s instructions. Analysis of each sample was performed in duplicate and run on two separate occasions. Cell lines were seeded to 10^6^ cells and after incubation for 24 h, the cells were counted and the culture supernatants (50 μL) were transferred in duplicate to plates precoated with cytokine-specific antibodies conjugated with different color-coded beads. The plates were then incubated for 1 hour, washed, and sequentially incubated with 50 μL of biotinylated cytokine-specific detection antibodies and a streptavidin-phycoerythrin conjugate. Positive and negative quality controls were included in duplicate in each assay. Fluorescence was recorded using a Bioplex 200 instrument. Cytokine/chemokine concentrations were calculated with Bio-Plex Manager software using a standard curve derived from recombinant cytokines. The results were normalized to million cells/mL.

### 2.8 MiRNA extraction, RT−PCR, and quantitative real-time PCR

miRNAs were extracted using the PureLinkTM miRNA Isolation Kit (Thermo Fisher Scientific). miRNA concentrations were determined using a QuBit 4 fluorometer and associated kits (Thermo Fisher Scientific).For cDNA synthesis of extracted miRNA from each cell line, RT−PCR was performed using the All-in-One miRNA First-Strand cDNA Synthesis Kit (GeneCopoeia) and 200 ng/μl of RNA per reaction. The real-time PCR assay was conducted in a CFXConnect Real-time PCR detection system (Bio-Rad). One microliter of cDNA was used per reaction, and PCRs were performed in quadruplicate. The values obtained are expressed as the means ± SDs. Genes with adjusted p values < 0.05 and fold changes > 2 were considered to be differentially expressed. qPCR of the miRNome was performed using the All-in-OneTM miRNA qRT−PCR Detection Kit 2.0 (GeneCopoeia). The PCR conditions included 1 cycle of 10 min at 95°C, followed by 40 cycles of 10 s of denaturation at 95°C, 30 s at 60°C, and 10 s at 72°C. The miProfileMouse miRNome miRNA qPCR Array (catalog QM002; GeneCopoeia) was used to analyze 834 mouse miRNAs.

To identify miRNAs potentially involved in the regulation of differentially expressed genes by the Dormant-met group, specific mouse miRNome databases, including miRDB and miRTarBase, were utilized. The miRDB is an online database for predicting functional microRNA targets (21). miRTarBase was also employed for comprehensive target prediction (22).

### 2.9 Isolation of lung leukocytes

Lungs were excised and collected 25 days after removal of the primary tumor. Each experimental group comprised 10 mice. The assays were conducted in duplicate. Lungs were dissociated into single-cell suspensions using a gentle MACS Dissociator (Miltenyi Biotech). Each lung was dissociated by loading a C tube on the machine with a volume of buffer (0.01 mol/L phosphate-buffered saline [PBS], 0.5% bovine serum albumin [BSA], and 2 mmol/L ethylenediaminetetraacetic acid [EDTA]). An installed program was selected for the dissociation process. After the completion of the program, whole-cell suspensions were centrifuged at 300 × g at room temperature for 10 min. The suspensions were collected and filtered through a 70-μm-pore-size nylon cell strainer to remove clumps and generate single-cell suspensions. Red blood cells were lysed with ACK lysis buffer (Gibco) for 5 min. The lysed cells were then washed twice in PBS. Viable cells were used for the antibody staining reaction.

### 2.10 Flow cytometry analysis of immune cell subsets

The following labeled antibodies from Miltenyi Biotec were utilized for the direct immunofluorescence study: CD45-PerCP, CD45-FITC, CD3-FITC, CD3-PE, CD19-PE, CD4-PerCP-Vio700, CD4-FITC-Viobright, CD8-PE, CD8-PerCP-Vio700, CD25FITC-Viobright, FoxP3-PE, CD49b-PE, MHC class II-PerCP-Vio700, CD11c-PE, CD11b-FITC-Viobright, F4/80-PE, Ly6G FITC, LY6C-FITC, Siglec-F-FITC, TCRαβ-PE, and TCRγδ-PerCP-Vio700. Isotype-matched nonimmune mouse IgGs conjugated with FITC, PE, or PerCP-Vio700 served as controls. FcR Blocking Reagent was used to block unwanted binding of antibodies to mouse cells expressing Fc receptors. Briefly, 5 × 10^5^ cells were washed twice with PBS. The cells were incubated for 10 min at 4°C in the dark with the primary antibodies. For Treg cell determination, cells were incubated for 10 min at 4°C with anti-CD4 and anti-CD25 antibodies after permeabilization for 30 min and incubated for 30 min at 4°C with anti-FoxP3 antibody. Leukocyte subpopulations were determined by gating total leukocytes based on their FSC/SSC profile and selecting CD45+ cells. The percentage of TCRγδ T lymphocytes was determined with respect to that of CD3+ T lymphocytes. The percentage of Treg cells was determined with respect to that of CD4+ T lymphocytes. The cells were analyzed on a FACSCanto cytometer (BD Bioscience). Each sample contained at least 5 × 10^4^ cells and was analyzed with CellQuest-Pro software.

### 2.11 Analysis of cell surface marker expression on metastatic tumor cells

Briefly, 5 × 10^5^ cells were washed twice with PBS and incubated for 10 min at 4°C with the following primary antibodies: anti-CD166-PE (Alcam), anti-CD95-PE (Fas), anti-VCAM1-PE, anti-CD21-FITC, anti-CD200R-FITC (Miltenyi-Biotech) and anti-CD276-FITC (Invitrogen). Isotype-matched nonimmune mouse IgG served as a control. A minimum of 1 × 10^4^ cells were analyzed with CellQuest Pro software.

### 2.12 Statistical analysis

Prism Software (Graph Pad Software V5.0) and SPSS Statistical 25.0 (IBM SPSS Inc., Chicago, IL, USA) were used for the statistical analyses. Two-tailed unpaired Student’s t-tests or Mann–Whitney U tests were used for statistical comparisons between two groups. ANOVA followed by the Tukey post hoc analysis or the Kruskal–Wallis test followed by the Dunn’s post hoc test was used for multiple comparisons. Fisher’s exact test was used to analyze the associations between two qualitative variables. Spearman rank correlation was applied to analyze the relationships between two variables. All the data are expressed as the means ± standard deviations (SD). p < 0.05 was considered to indicate statistical significance. Significance levels are denoted as follows: *p < 0.05,** p < 0.01,*** p < 0.001.

## 3 RESULTS

### 3.1 GR9-B11 dormant metastatic murine model

The GR9-B11 tumor cell line was derived from a fibrosarcoma induced by methylcholanthrene injection into BALB/c mice using limiting dilution cloning. Spontaneous metastasis assays were conducted using this tumor cell line in wild-type BALB/c mice, but no overt spontaneous metastasis was observed (Fig. 1A) (18). However, when the same assays were conducted in BALB/c nude mice, metastases were observed in 80% of the mice (Fig. 1A) (18). These metastases were referred to as Nude Metastases (Nude-met). Spontaneous metastasis assays were conducted in immunocompetent mice that were depleted of CD8+ T lymphocytes 5 months after primary tumor removal. The results showed that overt spontaneous metastases occurred in 100% of the mice (Fig. 1A) (18). The metastases remained dormant for five months in immunocompetent mice until T lymphocytes were depleted. They were then referred to as Dormant Metastasis (Dormant-met). Notably, the only action taken was the depletion of T lymphocytes in the host. No direct intervention was made on the dormant metastatic cells, yet this was sufficient for them to ‘awaken’ and begin to grow. Several metastases of each type were isolated and adapted to cell culture. For this study, we selected two nude metastases, Nude-met-1 and Nude-met-2, and two dormant metastases, Dormant-met-1 and Dormant-met-2 (Figure 1A). It is important to note that both metastasis groups originated from the same B11 tumor clone. One group was generated in immunodeficient BALB/c nude mice and their metastases had never experienced dormancy, while the other group was generated in immunocompetent mice depleted of T lymphocytes, and their metastases are in dormancy controlled by the immune system. To determine whether dormant metastasis retains their ability to remain dormant *in vivo*, we conducted spontaneous metastasis assays with Dormant Metastases in immunocompetent mice. This resulted in the generation of primary tumors, producing later metastasis in immunodormancy. (Fig. 1B). These recent assays demonstrated that Dormant Metastases retain their ability to enter a state of *in vivo* metastatic dormancy. In summary, the dormancy metastasis model we studied comprises the following cell lines: GR9-B11, which is a tumor clone; two Nude Metastases, namely, Nude-met1 and Nude-met2; and two Dormant Metastases, namely, Dormant-met1 and Dormant-met2 (Fig. 1A). This study also included two overt spontaneous metastases, Overt-met-1 and Overt-met-2, generated in wild-type BALB/c mice by another clone from the GR9 tumor system, GR9-A7 (Fig. 1A) (20). There are three groups of metastases: Dormant-met, Nude-met, and Overt-met.

**Figure 1.**
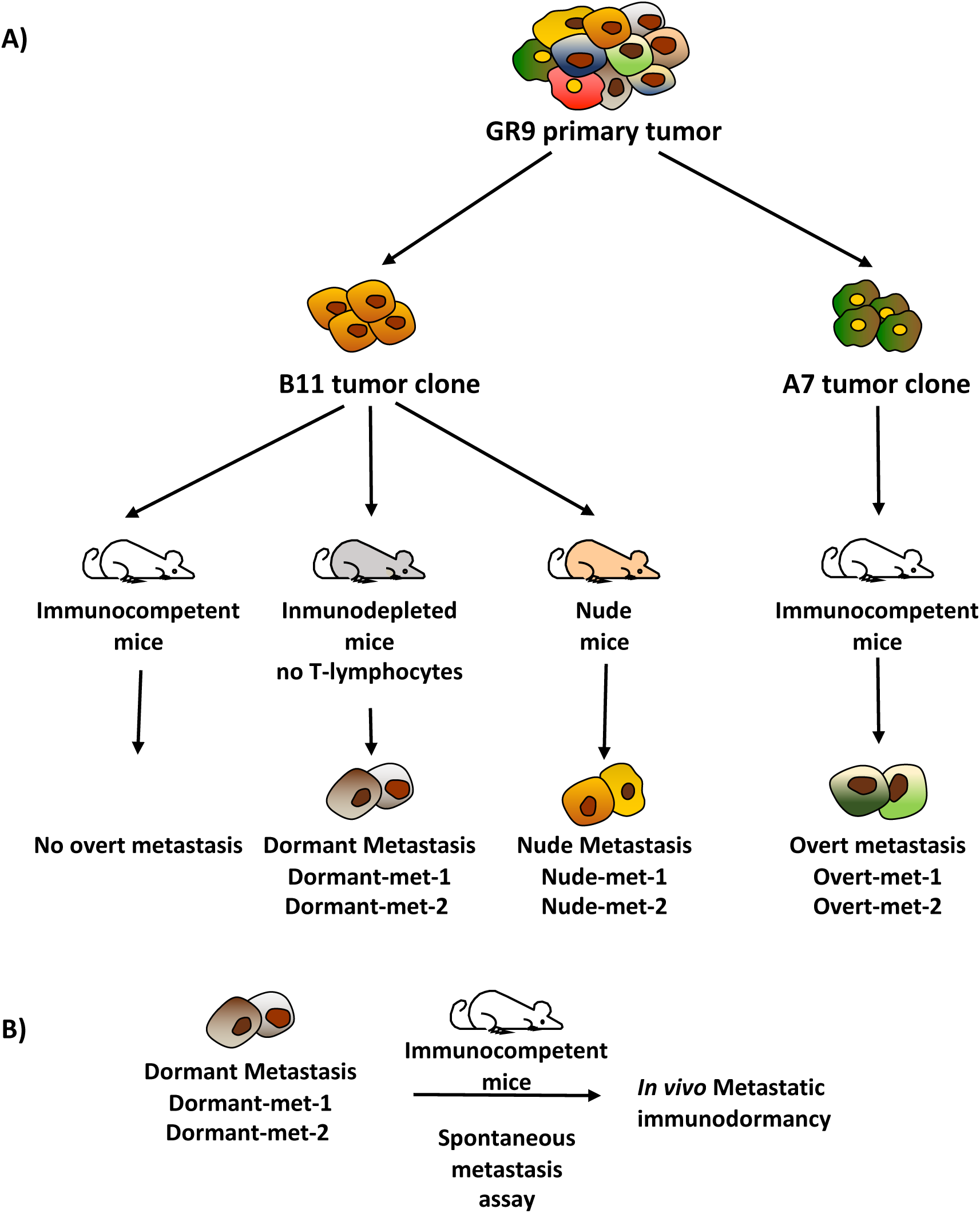
Murine tumor model of metastatic dormancy**. A,** Primary GR9 tumor was generated in BALB/c mice by injecting methylcholanthrene. This tumor was adapted for culture and cloning, and tumor clones such as B11 and A7 were obtained. In spontaneous metastasis assays in immunocompetent BALB/c mice, the B11 tumor clone did not generate overt spontaneous metastases. However, when T lymphocytes are depleted in these mice five months after removal of the primary tumor, the hosts develop spontaneous metastases, namely, Dormant-met-1 and Dormant-met-2. In nude BALB/c mice, B11 generates overt spontaneous metastases, namely, Nude-met-1 and Nude-met-2. The A7 tumor clone produces overt spontaneous metastases in immunocompetent BALB/c mice, specifically Over-met-1 and Overt-met-2. **B,** Spontaneous metastasis assays were performed by injecting Dormant Metastases into BALB/c mice, and immune-controlled dormant metastases were generated again.

### 3.2 Proliferation assays and treatment with chemotherapeutic agents and chemokines

As dormant metastases are exclusively in a state of dormancy, characterized by minimal or no proliferation, the initial assays aimed to determine whether there were differences in *in vitro* proliferation capacity among the different metastases. The *in vitro* growth of the various metastatic cell lines was found to be similar, with no significant differences observed in their growth patterns (Fig. 2A). The cells were subjected to nutrient-restricted conditions in culture by reducing 0.5% fetal bovine serum for 5 days. The proliferation rate of the cells was measured and compared to their proliferation rate under normal culture conditions with 10% fetal bovine serum. The results showed that dormant metastastatic cell lines exhibited greater resistance to these restrictive conditions, maintaining greater cell proliferation than did the nude and overt metastastatic cell lines (21.2% vs. 9.8% vs. 8.9%, respectively) (Fig. 2B). Under nutrient-restricted conditions, Dormant Metastases maintained a proliferation rate that was twice as high as that of the other two types of metastases. Metastases in dormancy may exhibit greater resistance to chemotherapeutics commonly used for anticancer treatments due to their low proliferation. Although they have equal *in vitro* proliferation rates, we measured their sensitivity to different chemotherapeutic agents commonly used in the clinic, including paclitaxel, cisplatin, methotrexate, and doxorubicin. The chemotherapeutic agents were used to treat the different metastases *in vitro*, and their proliferation rate was measured. The sensitivity of the detection of overt metastases was measured to determine a dose/response curve. Concentrations were selected based on the highest sensitivity exhibited by Overt-Met, which resulted in 100% inhibition of cell proliferation. Growth inhibition in the other two types of metastases was also measured. Nude Metastases exhibited inhibition values similar to those of Overt Metastases, with no significant differences (Fig. 2C). In contrast, the sensitivity of Dormant Metastases to paclitaxel and cisplatin was lower, exhibiting only 47% and 61% growth inhibition, respectively (Fig. 2C). However, compared with those of Overt-met and Nude-met, Dormant Metastases showed greater sensitivity to the other two chemotherapeutic agents, methotrexate and doxorubicin, resulting in 199% and 359% growth inhibition, respectively, compared to Overt-Met (Fig. 2C). The results suggest that Dormant Metastases have lower sensitivity to paclitaxel and cisplatin, two commonly used chemotherapeutics, despite having a proliferation rate similar to that of other metastases. In contrast, Dormant Metastases exhibit greater intrinsic sensitivity to methotrexate and doxorubicin.

**Figure 2.**
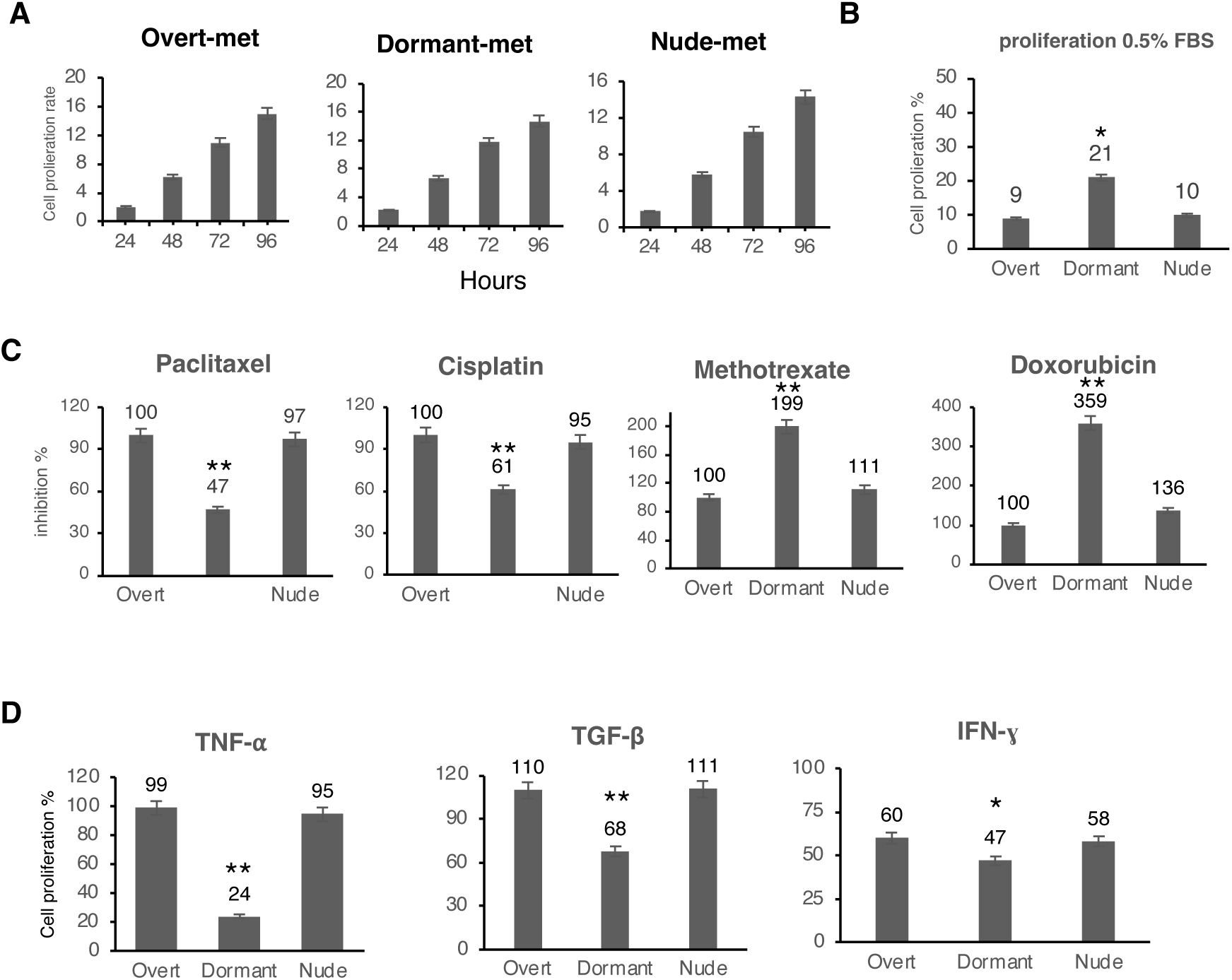
The *in vitro* proliferation of the three groups of metastases was studied under different conditions. **A,** The proliferation rate was measured under basal conditions. **B,** The proliferation rate was measured under nutrient-restricted conditions with 0.5% fetal bovine serum. **C,** Growth inhibition rate was measured after treatment with different chemotherapeutic agents: paclitaxel, cisplatin, methotrexate, and doxorubicin. **D,** The proliferation rate was measured after treatment with different cytokines: TNF-α, TGF-β, and IFN-γ. The values represent the means ± SDs of three independent experiments performed in triplicate. The statistical analysis was performed using ANOVA test, followed by Tukey’s post hoc test (*p < 0.05; **p < 0.01).

In other experiments, cells were exposed to TNF-α, TGF-β, and IFN-γ, which are all involved in the immune response, to assess their effect on *in vitro* proliferation. The proliferation of cells without any treatment was considered 100%, and the percentage of proliferation resulting from different treatments was measured. The results indicated that treatment with TNF-α did not significantly inhibit growth in Overt-met or Nude-met, with proliferation rates of 99% and 95%, respectively. However, it strongly inhibited growth in Dormant-met, with a proliferation rate of only 24% (Fig. 2D). Treatment with TGF-β had a slight stimulatory effect on the growth of Overt Metastases and Nude Metastases, with proliferation rates of 110% and 111%, respectively. In contrast, it significantly inhibited the growth of Dormant Metastases, with a proliferation rate of 68% (Fig. 2D). Treatment with IFN-γ inhibited growth in Overt and Nude Metastases by 60% and 58%, respectively. Additionally, IFN-γ significantly inhibited the growth of Dormant Metastases by 47% (Fig. 2D). In summary, Dormant-met exhibited greater *in vitro* growth inhibition when treated with all three chemokines, particularly TNF-α, suggesting a greater sensitivity to these treatments.

### 3.3 Comparison of transcriptional gene expression among different metastasis groups

mRNA sequencing was performed to compare differential gene expression at the transcriptional level. The study analyzed the gene expression of different metastasis groups, each consisting of two metastases, including the GR9-B11 tumor clone. The Pearson correlation coefficient of gene expression between the two metastases in each group was R² > 0.975. Additionally, the differential expression of Dormant-met group was compared to that in the other two metastasis groups and the GR9-B11 tumor clone group, and the expression in these latter cells was used as a control. Ninety-two genes with differential expression were found in the Dormant-met group (Fig. 3 and Supplemental Table S1). In the Dormant-met group, 65 genes exhibited overexpression, while 27 genes demonstrated lower expression compared to the other two metastasis groups and the B11 clone (Fig. 3). This set of 92 genes exhibited a phenotype of differential gene expression at the transcriptional level for the Dormant-met group. Notably, if a gene exhibited greater expression in the Dormant-met group, it maintains this greater expression when compared to Nude-met and Overt-met. Similarly, if the expression is lower in the Dormant-met group, it remains with lower expression when compared to the other two metastasis groups. The expression of the Rspo1 gene showed the most significant difference between the Dormant-Met group and the other two metastasis groups, exhibiting very high expression in the Dormant-met group. Analysis of B11 tumor clone expression revealed that the expression of certain genes increased in the Dormant-Met group and decreased in the Nude-met group. These genes included Ch25h, Alcam, Fas, Ccl2, and Ccl7.

**Figure 3.**
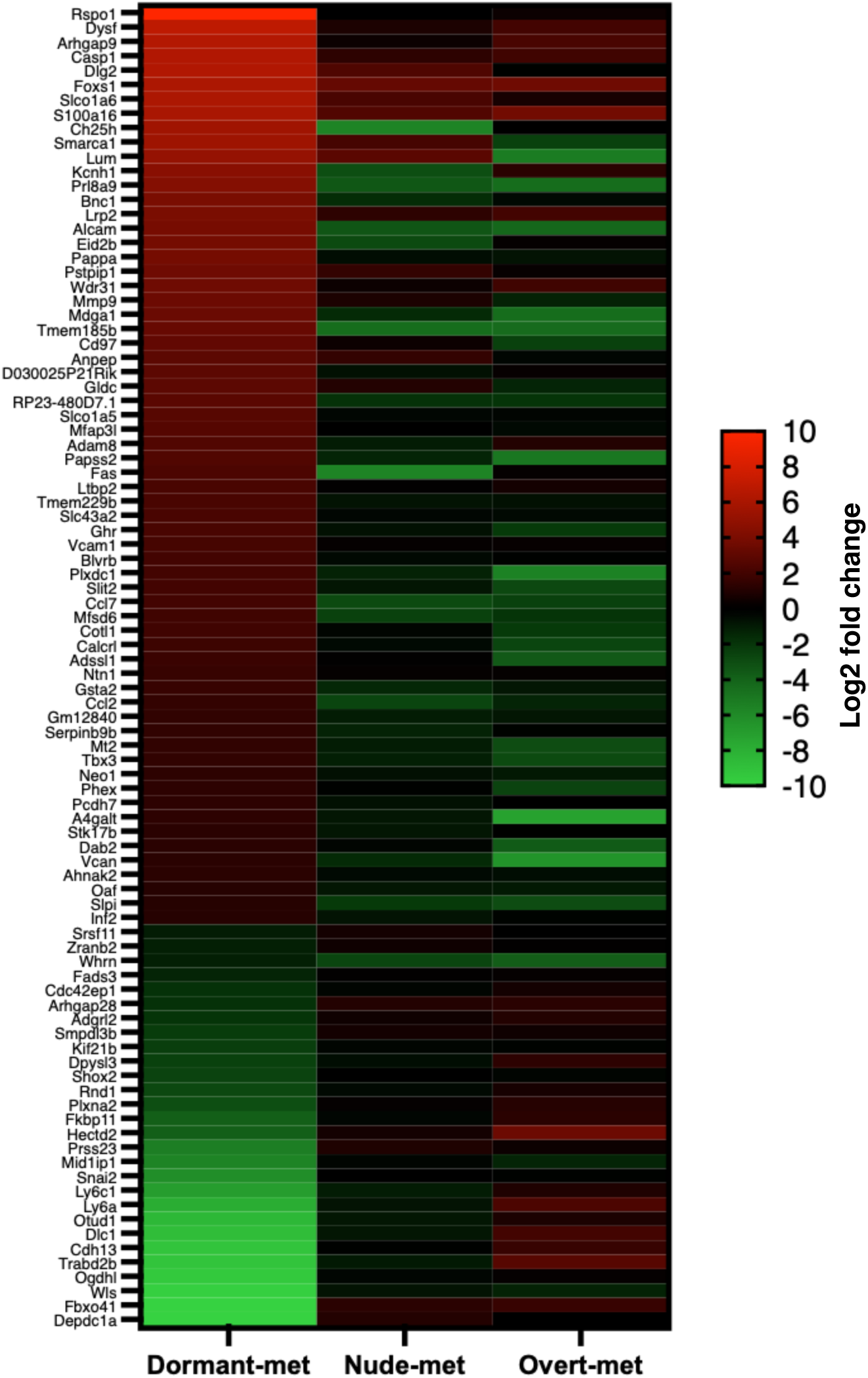
Genes differentially expressed in the Dormant Metastasis group. A heatmap is presented to illustrate the differential expression data between the three metastatic groups obtained from RNA-Seq. Only genes that displayed a greater than twofold difference in expression (log2>1) with a p value less than 0.001 were identified and plotted in the heatmap. The expression values are presented as log2 (counts per million) values and are scaled by gene.

Further analysis of these results through real-time qPCR plates was performed to examine the expression of genes associated with the cholesterol pathway, surface markers, and chemokines. The metastasis groups were compared, and relative expression was calculated by using Overt-met as a control and assigning it a value of 1. Only differences with a value greater than 2 and significant (p < 0.05) were considered. Compared with that of Nude-met group, the mRNA sequencing of Dorm-met group showed the greatest increase in expression of the Ch25h enzyme (log2: 11.3). Furthermore, there was an increase in Dormant (log2: 5.6) and a decrease in the Nude group (−5.7) compared with the Overt-met data (log2: 0). This enzyme is responsible for converting cholesterol into 25-hydroxycholesterol. Therefore, we measured the expression of genes involved in the cholesterol pathway. The expression of 13 genes significantly increased in the Dormant-Met group compared to the other two metastasis groups: Mvd, Pmvk, Nshdl, Tm7sf2, Hmgcr, Cyp51, Srebf2, Idi1, Dhcr7, Hmgcs1, Fdft1, Ebp, and Mvk (Fig. 4A). This finding suggested that there may be an increase in cholesterol synthesis in Dormant-Met, which is then converted into 25-hydroxycholesterol.

**Figure 4.**
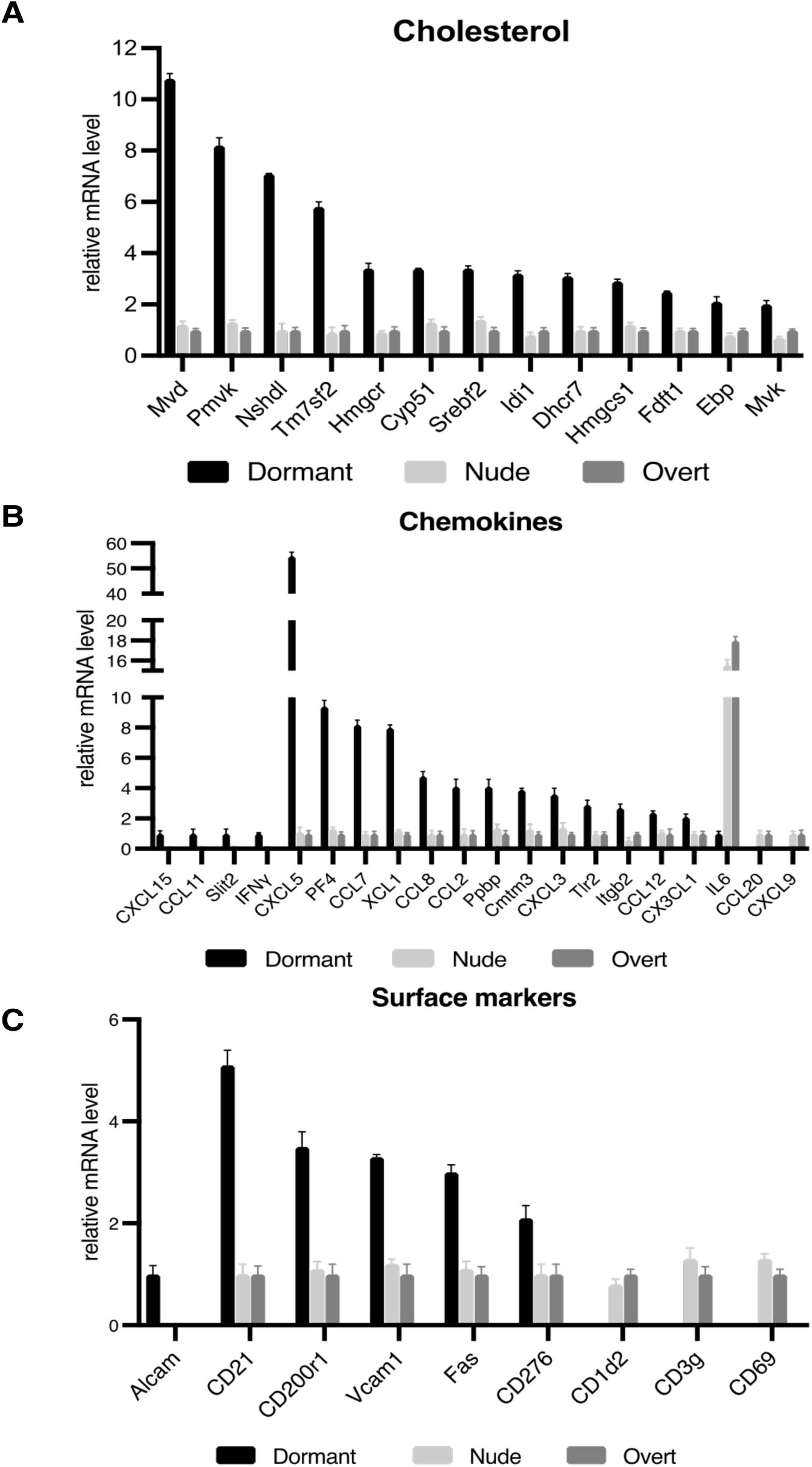
Analysis of differentially expressed genes in the Dormant Metastasis group was performed by RT−qPCR. **A,** Genes involved in the cholesterol synthesis pathway. **B,** Chemokine genes. **C,** Surface marker genes. Genes that showed more than a 2-fold difference (p < 0.01) when comparing the Dormant-met with Nude-met and with Overt-met were plotted. The expression levels of the genes of interest were determined with respect to the levels of the β-actin and GAPDH housekeeping genes. The data for the Overt-met group were set to 1. The values represent the means ± SDs of three independent experiments performed in duplicate. The statistical analysis was performed using ANOVA, followed by the Tukey’s post hoc test.

Analysis of chemokines revealed that only Dormant-met expressed Cxcl15, Ccl11, Slit2, and IFN-γ (Fig 4B). The results from Dormant-met showed a significant increase in Cxcl5 and PF4, Xcl1, Ccl8, Pbpb, Cmtm3, Cxcl3, Tlr2, Itgb2, Ccl12, and Cx3cl1. Additionally, the study confirmed the increased expression of Ccl7 and Ccl2. In contrast, Dormant-met showed a considerable decrease in IL-6 expression and a complete absence of Cxcl9 and Ccl20 (Fig. 4B).

There was an increase in the expression of six surface markers (Fig. 4C). This finding supports the RNA-seq data for Alcam, which is exclusively expressed in Dormant-met, and for Fas and Vcam1, which exhibited a significant increase. Furthermore, the expression of three additional markers increased: CD276 (B7-H3), CD21 (Cr2), and CD200r1. All of these markers are involved in interactions with immune cells. In contrast, Dormant-met showed a lack of expression of with immune cells. In contrast, Dormant-met showed a lack of expression of CD1d2, CD3g, and CD69, whereas the other two metastasis groups exhibited expression (Fig. 4C). According to the flow cytometry assays, among three metastasis group, only the Dormant-Met group expressed Alcam on the cell surface among the three metastasis groups (Fig. 5). The surface expression of other markers remained unchanged.

**Figure 5.**
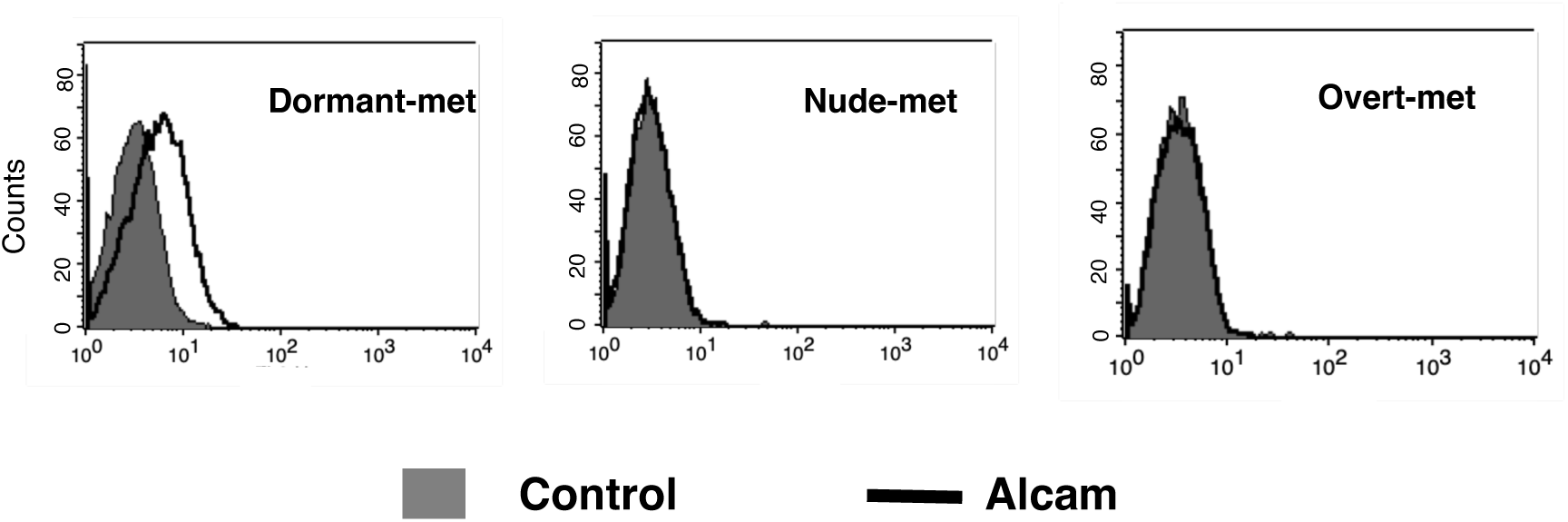
The Alcam surface expression on the three metastatic groups. Flow cytometry analysis showed that Alcam protein is expressed exclusively in the Dormant-met group.

### 3.4 Chemokines secreted by various metastases

The analysis focused on the cytokines secreted by different metastasis groups into the culture medium, which could affect immune cells. This study revealed a noteworthy increase in the secretion of the following chemokines by Dormant-met: Ccl2, Ccl7, Cxcl1, Cxcl2, Cxcl5, TGF-B1, Ccl8, and TGF-B2. Additionally, unlike the other two metastasis groups, the Dormant-met group stopped secreting IL6. (Fig. 6).

**Figure 6.**
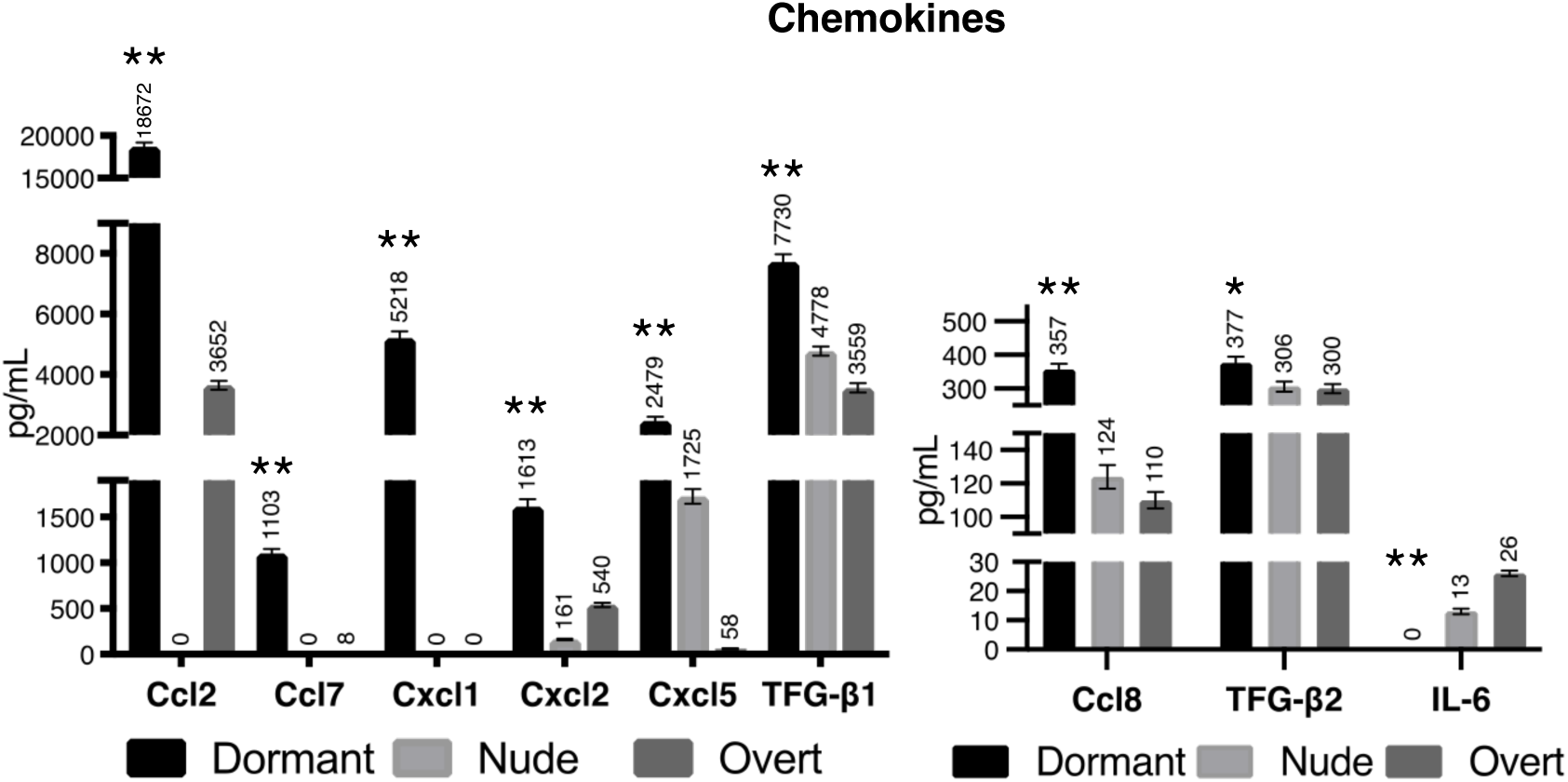
Cytokine/chemokine profile of the Dormant Metastasis group versus the Nude-met and Overt-met groups. Only cytokines/chemokines whose expression changed are depicted. The values are presented as the means ± SDs from two independent experiments performed in duplicate. The statistical analysis was performed using ANOVA test, followed by Tukey’s post hoc test (*p < 0.05; **p < 0.01).

### 3.5 Differentially expressed miRNAs in Dormant Metastases

MicroRNAs are important regulators of gene expression within cells. the mouse miRNome, which comprises 834 miRNAs was analyzed for each metastasis group. The expression of miRNAs in Dormant-Met or Nude-Met was examined using Overt-met as a control with a value of 1. This study revealed that nine miRNAs had significantly increased expression exclusively in the Dormant-met group compared to the other two metastasis groups. These miRNAs are miR-142-3p, miR-380-5p, miR-34b-3p, miR-376b-3p, miR-21-5p, miR25-3p, miR-1930-5p, miR-5046, and miR-7a-5p. (Fig. 7A). It is important to note that miR-142-3p is expressed exclusively in Dormant-met. In contrast, 11 miRNAs were weakly expressed only in Dormant-met, exhibiting a significant decrease in expression compared to the other two metastasis groups. These miRNAs are miR-382-3p, miR-300-5p, miR-669o-5p, miR-3060-5p, miR-3079-3p, miR-21-3p, miR-211-5p, mR-3094-3p, miR-499-3p, miR-154-3p, and miR-467d-5p. (Fig. 7B). Notably, miR-467d-5p was not expressed in the Dormant-met group, but was expressed in the other two metastasis groups.

We analyzed miRNA databases to identify miRNAs that may contribute to the differential gene expression observed in the Dormant-met group. A reduction in miR-211-5p expression may result in increased expression of five genes: Cxcl5, Alcam, Mfsd6, Lrp2, and Stk17b. A decrease in miR-21-3p expression may lead to increased expression of three genes: Insig2, Cxcl15, and Neo1. The upregulation of Cxcl15 may also be mediated by the reduced expression of miR-382-3p. Consequently, miRNAs tightly regulate the expression of chemokines. This was also observed for Ccl2 and Ccl7, whose increased expression in Dormant-met group could be regulated by the loss of expression of miR-499-3p and miR-154-3p, respectively. During the dormant metastasis process, decreased miR-3094-3p expression may lead to increased Ch25h gene expression. Conversely, increased miR-142-3p expression may cause decreased IL-6 and Hectd2 gene expression.

**Figure 7.**
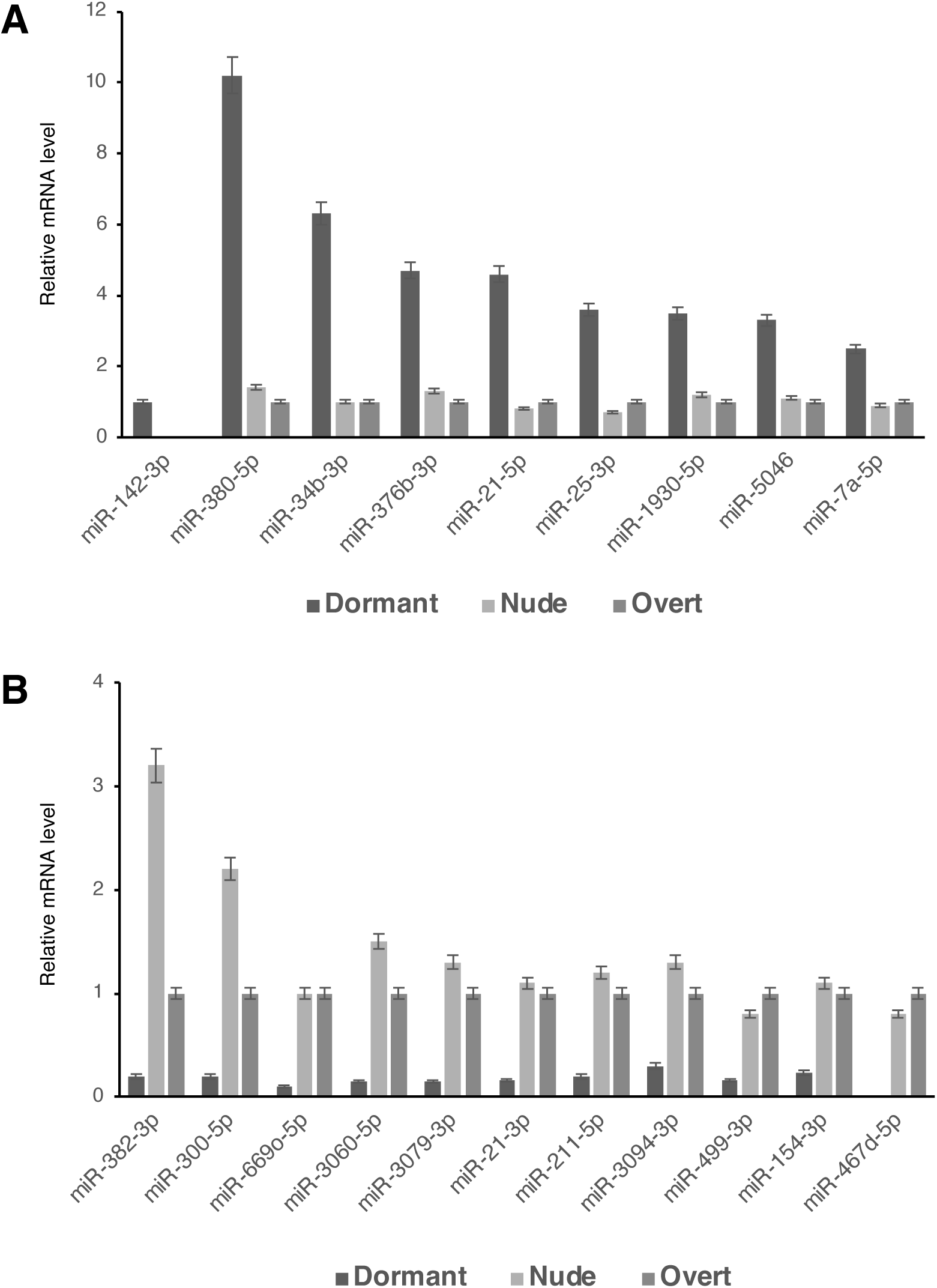
MicroRNAs that are differentially expressed in the Dormant Metastasis group. **A,** miRNAs that are upregulated in the Dormant-met group. **B,** miRNAs whose expression was downregulated in the Dormant-met group. The expression of 834 miRNAs was measured using the ‘mouse miRNome qPCR Arrays 18.0’ (Genecopoeia). Only miRNAs that displayed a greater than twofold difference (p < 0.01) when comparing the Dormant-met group with the Nude-met and Overt-met groups were plotted. The expression levels of the miRNAs of interest were determined with respect to the levels of six housekeeping snRNAs (MK1-MK6). The data for the Overt-met group were set to 1. The values are presented as the means ± SDs of three independent experiments, each performed in duplicate. The statistical analysis was conducted using ANOVA test, followed by a Tukey’s post hoc test.

### 3.6 Immune cells in the microenvironment of Dormant Metastases

In the microenvironment of dormant metastases, an immune response may have originated that could control the metastases, although they cannot be destroyed. We analyzed various immune subpopulations in the lungs of animals with Dormant Metastases, 25 days after the removal of the primary tumor, and compared them with those of wild-type animals (control). In mice injected with B11, there was a significant increase in T lymphocytes (64.5% vs. 51.9%), including both CD4+ helper T lymphocytes (46.2% vs. 40.6%) and CD8+ cytotoxic T lymphocytes (17.4% vs. 9.7%) (Fig. 8). These cells may be involved in the immune control of dormant metastases. In addition, the results showed a significant increase in neutrophils (9.4% vs. 4.6%) and γδ T cells (13.5% vs. 7.4%) in mice injected with B11 (Fig. 8). No significant differences were detected in the other analyzed populations.

**Figure 8.**
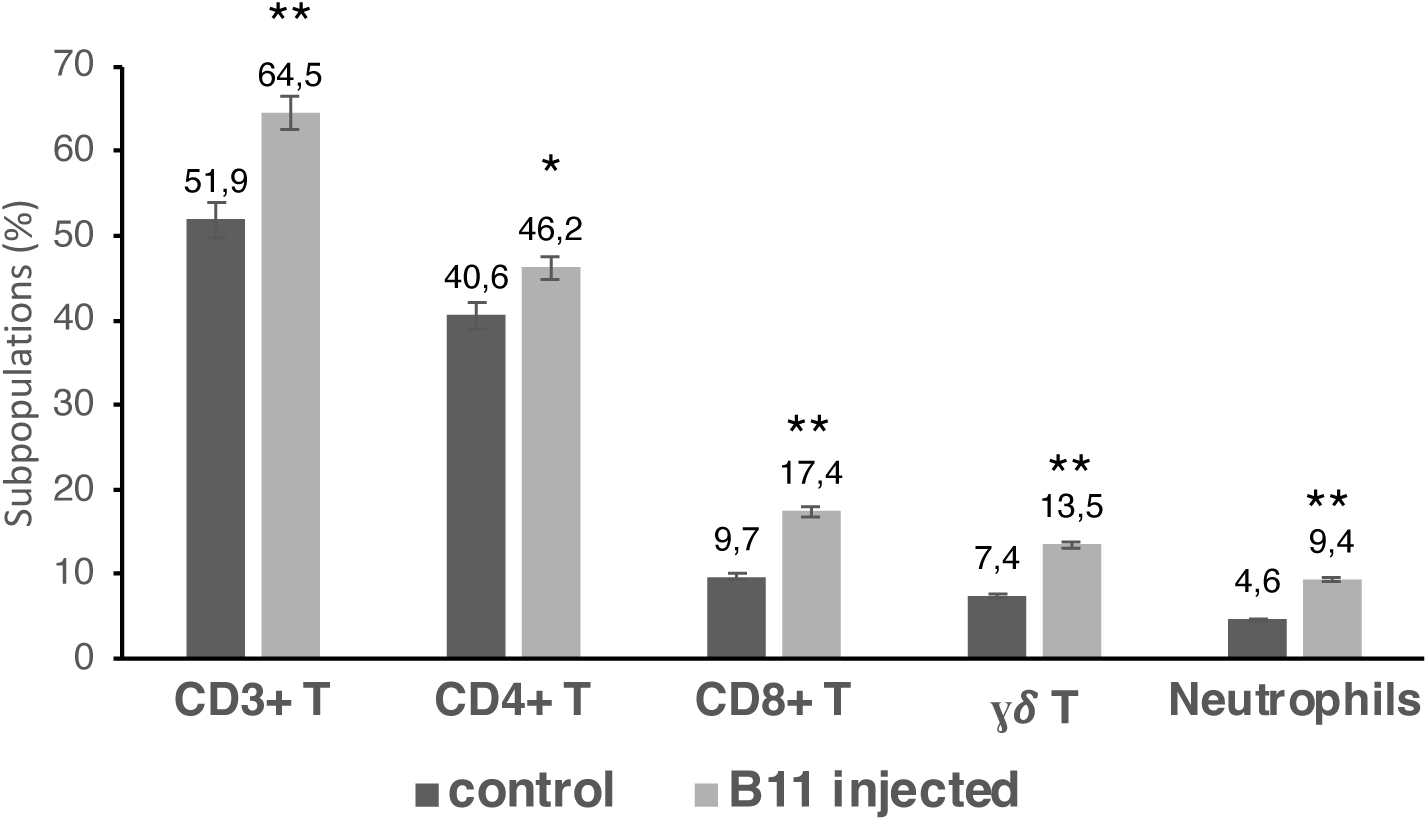
Changes in lung leukocyte populations. On day 25 after local tumor removal, we analyzed the various leukocyte populations by comparing B11 tumor-injected mice (B11-injected) with nontumor-injected mice (control). Only leukocyte populations that exhibited significant changes in their expression are presented. The data are expressed as the means ± SDs of 20 mice in each group. *p < 0.05; **p < 0.01. The statistical analysis was performed using a two-tailed Student’s t test.

## 4 Discussion

Metastatic dormancy is a crucial stage in cancer progression. At this stage, metastatic cells cease to proliferate and enter a state of quiescence or very low proliferation (6). They cannot be entirely eliminated by the immune system and may later develop into overt metastases. Although this process is poorly understood, several groups have recently investigated how these dormant metastatic cells exit this dormant state to resume proliferation (23–26). This study examined dormant metastases before they could escape, aiming to identify their phenotype and the immune cells involved in the dormant metastatic state. To achieve this goal, the study did not directly awaken dormant metastatic cells. Instead, depletion of the host immune system allows the cells to proliferate and progress, resulting in overt metastases. Furthermore, these dormant metastatic cells did not enter a state of quiescence per se, as they exhibited the same *in vitro* proliferation capacity as did overt metastases. Conversely, when the different metastatic cell lines were subjected to nutrient restriction, the Dormant Metastases group exhibited increased proliferation compared to the Overt and Nude Metastases group. Moreover, when these dormant metastatic cells were treated *in vitro* with chemotherapeutic agents commonly used in clinical therapy, varying levels of sensitivity were observed compared to cell lines derived from Overt Metastases (Overt-met) or from Nude Metastases (Nude-met). Cell lines derived from Dormant Metastases (Dormant-met) showed decreased sensitivity to paclitaxel and cisplatin but higher sensitivity to methotrexate and doxorubicin. These findings suggest that although these cells have a similar *in vitro* proliferation capacity to that of overt metastases, they still inherently maintain greater resistance to specific chemotherapeutics such as paclitaxel and cisplatin. Reactivating the cell cycle in dormant metastases to to increase their sensitivity to these chemotherapeutics may not be effective (27). However, these results suggest that treatments such as methotrexate and doxorubicin can halt proliferation and even destroy to dormant metastases. Furthermore, it was observed that cell lines derived from Dormant Metastases exhibit greater inhibition of proliferation when treated *in vitro* with the chemokines TNF-α, TGF-β, and IFN-γ. This finding suggested that the secretion of these cytokines by immune system cells could induce a nonproliferative state in these dormant metastatic cells *in vivo*. Additionally, these chemokines could be used to treat patients and maintain a nonproliferative state of dormant metastases.

The study’s findings indicate that dormant metastatic cells can re-enter metastatic dormancy *in vivo* even when an intact host immune system is present. The immune system is responsible for maintaining immune-mediated dormancy in these cells *in vivo*. Notably, in our model, Dormant Metastases expressed MHC-I molecules on their surface, despite originating from an MHC-I-negative tumor clone (18). This, along with the increase in CD3 T lymphocytes in the lungs, may enable the control of these cells, although complete destruction may not be possible. There is also an increase in neutrophils and γδ T lymphocytes, which may have an immunosuppressive effect on CD3 T lymphocytes (28–32).

Analysis of gene expression through massive mRNA sequencing revealed that dormant metastasi group exhibited a distinct pattern. In this metastasis group, 92 genes were differentially expressed compared to Nude-met and Overt-met groups. This specific phenotype for Dormant Metastases is defined by this gene expression pattern, with 65 genes exhibiting increased expression. In Dormant-met, one gene that is highly expressed is Ch25h, an enzyme responsible for converting cholesterol into 25-hydroxycholesterol. This oxysterol can bind to LXR receptors and activate the transcription of genes involved in cholesterol flux (33). Thirteen highly expressed genes were found to be involved in cholesterol synthesis pathways. Our dormant metastatic cells increase cholesterol synthesis to transform it into 25-hydroxycholesterol. This molecule has been implicated in immunosuppressive processes, such as reducing antigen processing by dendritic cells and inhibiting their migration to lymphoid organs (34). Recently, it has been demonstrated that extracellular 25-hydroxycholesterol suppresses cholesterol biosynthesis in activated and proliferating T cells, inhibiting their cellular growth in a paracrine manner (35). Furthermore, it has been demonstrated that oxysterols promote the recruitment of neutrophils that support tumor growth (36). Our results indicate a greater concentration of neutrophils in the lungs of mice with dormant metastases. Additionally, 25-hydroxycholesterol has been linked to the maintenance and activation of γδ T cells (37). Our findings indicate an increase in γδ T cells in the lungs of mice with dormant metastases. These cells, along with neutrophils, may have an immunosuppressive role (31, 32). These results suggest that dormant metastatic cells may prevent their destruction by CD8+ T lymphocytes through increased Ch25h expression and 25-hydroxycholesterol production. CD8+ T lymphocytes are found in greater proportions in the lungs of mice with dormant metastases. In addition, dormant metastatic cells produce chemokines such as Ccl2, Ccl7, Cxcl1, Cxcl2, and Cxcl5. These chemokines can attract myeloid-derived suppressor cells (38–40).

Dormant metastasis group cells exhibited a distinctive positive transcriptional expression of the Alcam molecule, which results in a unique surface expression on dormant metastatic cells. Alcam is expressed in human tumors and can serve as a prognostic factor and metastasis predictor (41). Furthermore, the interaction of CD166 with the CD6 receptor on γδ T cells has been implicated in their activation (42). During the dormancy process in our model, Alcam may perform this function by suppressing cytotoxic T cells. Rspo1 is a soluble ligand that can transform M1 macrophages into an immunosuppressive M2 phenotype, blocking CD8+ T lymphocytes (43). This molecule may play an immunosuppressive role in our model. In summary, an increase in T lymphocytes may maintain dormant metastatic cells in a state of dormancy. However, immunosuppressive mechanisms promoted by dormant metastatic cells themselves prevent complete destruction by cytotoxic T lymphocytes. It is widely acknowledged that the immunosuppressive mechanisms mentioned above contribute to the evasion of tumor cells from the immune system. However, in this context, we focused on their role in the dormant metastatic process, prior to their ability to evade the immune system. These results indicate that inhibiting these immunosuppressive mechanisms during the dormant metastatic state might result in the eradication of dormant metastases in patients.

Notably, miR-142-3p was exclusively expressed in the dormant metastasis group. MiR-142 is a microRNA that is predominantly found in hematopoietic cells (44). It plays an important role in the immune system, specifically in Treg cells and macrophages (45, 46). MiR-142 has an anti-inflammatory role by downregulating IL-6, as observed in our dormant metastases (47, 48). In the dormancy phase, the expression of miR-142-3p is significantly increased during the latent phase of tuberculosis infection compared to the active phase (49). This results in a decrease in IL-6 expression due to the downregulation of IRAK-1 (50).

Our study revealed that Dormant Metastases have a unique and specific phenotype that differentiates them from Overt-met or Nude-met, which are not in a state of dormancy. This phenotype has various characteristics, including different sensitivities to restrictive nutrient conditions, chemotherapeutic agents, and cytokines; differential expression of genes and miRNAs; distinct expression of surface markers; secretion of different chemokines; and the coexistence of T lymphocytes and immunosuppressive cells in the dormant metastatic microenvironment. The characteristics mentioned above may help to develop new biomarkers for dormant metastatic disease in patients where clinically undetectable dormant metastases are present. Tumors can be evaluated by measuring the expression of various genes or miRs, such as Ch25h, Alcam, Rspo1, and miR-142-3p. Alternatively, liquid biopsy can detect increased expression of chemokines such as Ccl2, Ccl7, Cxcl1, Cxcl2, and Cxcl5, or of molecules such as 25-hydroxycholesterol and Rspo-1, as well as miRNA 142-3p. Even a rise in cholesterol levels in patients might indicate the presence of dormant metastases. Additionally, studying the different immune cell populations in patients, both in the tumor and peripheral blood, could be useful for detecting increases in T lymphocytes and immunosuppressive neutrophils or γδ T cells.

## Declarations

### Ethics approval

This study was carried out in accordance with the recommendations of the European Community Directive 2010/63/EU and Spanish law (Real Decreto 53/2013) on the use of laboratory animals, and their housing and the experimental procedures were approved by the Junta de Andalucía animal care committee and adhered to the animal welfare guidelines of the National Committee for Animal Experiments

### Availability of data and materials

The data supporting the fundings of this paper is included in this paper and its supplementary file. The datasets used and /or analyzed in this study are available from the corresponding author on reasonable request.

### Authors’ contributions

IA and AMGL: design and supervision the study, wrote and revised of the manuscript. VC, IA, VS, MP, IR, EC, MM, ME, IL and AMGL: Development of methodology, acquisition of data and analysis and interpretation of data. All the authors revised the work, approved the final version, and agreed to be accountable of the work.

## Acknowledgements

Not applicable

## Supplementary information

### Additional File 1

Table S1.pdf. Differentially expressed genes in Dormant metastases obtained by RNA-seq.

## References

1. Mani K, Deng D, Lin C, Wang M, Hsu ML, Zaorsky NG. Causes of death among people living with metastatic cancer. Nat Commun. 2024;15(1):1519.

2. Massagué J, Obenauf AC. Metastatic colonization by circulating tumour cells. Nature. 2016;529(7586):298–306.

3. Fares J, Fares MY, Khachfe HH, Salhab HA, Fares Y. Molecular principles of metastasis: a hallmark of cancer revisited. Signal Transduct Target Ther. 2020;5(1):28.

4. Park SY, Nam JS. The force awakens: metastatic dormant cancer cells. Exp Mol Med. 2020;52(4):569–81.

5. Phan TG, Croucher PI. The dormant cancer cell life cycle. Nat Rev Cancer. 2020;20(7):398–411.

6. Truskowski K, Amend SR, Pienta KJ. Dormant cancer cells: programmed quiescence, senescence, or both? Cancer Metastasis Rev. 2023;42(1):37–47.

7. Dittmer J. Mechanisms governing metastatic dormancy in breast cancer. Semin Cancer Biol. 2017;44:72–82.

8. Cackowski FC, Heath EI. Prostate cancer dormancy and recurrence. Cancer Lett. 2022;524:103–8.

9. Gomis RR, Gawrzak S. Tumor cell dormancy. Mol Oncol. 2017;11(1):62–78.

10. Risson E, Nobre AR, Maguer-Satta V, Aguirre-Ghiso JA. The current paradigm and challenges ahead for the dormancy of disseminated tumor cells. Nat Cancer. 2020;1(7):672–80.

11. Wikman H, Vessella R, Pantel K. Cancer micrometastasis and tumour dormancy. APMIS. 2008;116(7-8):754–70.

12. Goddard ET, Bozic I, Riddell SR, Ghajar CM. Dormant tumour cells, their niches and the influence of immunity. Nat Cell Biol. 2018;20(11):1240–9.

13. Romero I, Garrido F, Garcia-Lora AM. Metastases in immune-mediated dormancy: A new opportunity for targeting cancer. Cancer Research. 2014;74(23):6750–7.

14. Sosa MS, Bragado P, Aguirre-Ghiso JA. Mechanisms of disseminated cancer cell dormancy: an awakening field. Nat Rev Cancer. 2014;14(9):611–22.

15. Aguirre-Ghiso JA. Translating the Science of Cancer Dormancy to the Clinic. Cancer Res. 2021;81(18):4673–5.

16. Gu Y, Bui T, Muller WJ. Exploiting Mouse Models to Recapitulate Clinical Tumor Dormancy and Recurrence in Breast Cancer. Endocrinology. 2022;163(6).

17. Bushnell GG, Deshmukh AP, den Hollander P, Luo M, Soundararajan R, Jia D, et al. Breast cancer dormancy: need for clinically relevant models to address current gaps in knowledge. NPJ Breast Cancer. 2021;7(1):66.

18. Romero I, Garrido C, Algarra I, Collado A, Garrido F, Garcia-Lora AM. T lymphocytes restrain spontaneous metastases in permanent dormancy. Cancer Res. 2014;74(7):1958–68.

19. Garrido A, Pérez M, Delgado C, Garrido ML, Rojano J, Algarra I, et al. Influence of class I H-2 gene expression on local tumor growth. Description of a model obtained from clones derived from a solid BALB/c tumor Intratumor Heterogeneity: Evolution through Space and Time Inverse relationship of melanocyte differentiation antigen expression in melanoma tissues and CD8+ cytotoxic-T-cell responses: evidence for immunoselection of antigen-loss variants in vivo. Experimental and clinical immunogenetics. 1986;3(2):98–110.

20. Romero I, Garrido C, Algarra I, Chamorro V, Collado A, Garrido F, et al. MHC Intratumoral Heterogeneity May Predict Cancer Progression and Response to Immunotherapy. Front Immunol. 2018;9:102.

21. Chen Y, Wang X. miRDB: an online database for prediction of functional microRNA targets. Nucleic Acids Res. 2020;48(D1):D127–D31.

22. Huang HY, Lin YC, Cui S, Huang Y, Tang Y, Xu J, et al. miRTarBase update 2022: an informative resource for experimentally validated miRNA-target interactions. Nucleic Acids Res. 2022;50(D1):D222–D30.

23. Albrengues J, Shields MA, Ng D, Park CG, Ambrico A, Poindexter ME, et al. Neutrophil extracellular traps produced during inflammation awaken dormant cancer cells in mice. Science. 2018;361(6409).

24. Fane ME, Chhabra Y, Alicea GM, Maranto DA, Douglass SM, Webster MR, et al. Stromal changes in the aged lung induce an emergence from melanoma dormancy. Nature. 2022;606(7913):396–405.

25. Hu J, Sánchez-Rivera FJ, Wang Z, Johnson GN, Ho YJ, Ganesh K, et al. STING inhibits the reactivation of dormant metastasis in lung adenocarcinoma. Nature. 2023;616(7958):806–13.

26. Sun D, Singh DK, Carcamo S, Filipescu D, Khalil B, Huang X, et al. MacroH2A impedes metastatic growth by enforcing a discrete dormancy program in disseminated cancer cells. Sci Adv. 2022;8(48):eabo0876.

27. Recasens A, Munoz L. Targeting Cancer Cell Dormancy. Trends Pharmacol Sci. 2019;40(2):128–41.

28. Germann M, Zangger N, Sauvain MO, Sempoux C, Bowler AD, Wirapati P, et al. Neutrophils suppress tumor-infiltrating T cells in colon cancer via matrix metalloproteinase-mediated activation of TGFβ. EMBO Mol Med. 2020;12(1):e10681.

29. Wculek SK, Malanchi I. Neutrophils support lung colonization of metastasis-initiating breast cancer cells. Nature. 2015;528(7582):413-7.

30. Emmons TR, Giridharan T, Singel KL, Khan ANH, Ricciuti J, Howard K, et al. Mechanisms Driving Neutrophil-Induced T-cell Immunoparalysis in Ovarian Cancer. Cancer Immunol Res. 2021;9(7):790–810.

31. Chabab G, Barjon C, Bonnefoy N, Lafont V. Pro-tumor γδ T Cells in Human Cancer: Polarization, Mechanisms of Action, and Implications for Therapy. Front Immunol. 2020;11:2186.

32. Fleming C, Morrissey S, Cai Y, Yan J. γδ T Cells: Unexpected Regulators of Cancer Development and Progression. Trends Cancer. 2017;3(8):561–70.

33. Calkin AC, Tontonoz P. Transcriptional integration of metabolism by the nuclear sterol-activated receptors LXR and FXR. Nat Rev Mol Cell Biol. 2012;13(4):213–24.

34. Villablanca EJ, Raccosta L, Zhou D, Fontana R, Maggioni D, Negro A, et al. Tumor-mediated liver X receptor-alpha activation inhibits CC chemokine receptor-7 expression on dendritic cells and dampens antitumor responses. Nat Med. 2010;16(1):98–105.

35. Takahashi H, Nomura H, Iriki H, Kubo A, Isami K, Mikami Y, et al. Cholesterol 25-hydroxylase is a metabolic switch to constrain T cell-mediated inflammation in the skin. Sci Immunol. 2021;6(64):eabb6444.

36. Raccosta L, Fontana R, Maggioni D, Lanterna C, Villablanca EJ, Paniccia A, et al. The oxysterol-CXCR2 axis plays a key role in the recruitment of tumor-promoting neutrophils. J Exp Med. 2013;210(9):1711–28.

37. Frascoli M, Ferraj E, Miu B, Malin J, Spidale NA, Cowan J, et al. Skin γδ T cell inflammatory responses are hardwired in the thymus by oxysterol sensing via GPR183 and calibrated by dietary cholesterol. Immunity. 2023;56(3):562–75.e6.

38. Teijeira Á, Garasa S, Gato M, Alfaro C, Migueliz I, Cirella A, et al. CXCR1 and CXCR2 Chemokine Receptor Agonists Produced by Tumors Induce Neutrophil Extracellular Traps that Interfere with Immune Cytotoxicity. Immunity. 2020;52(5):856–71.e8.

39. Takacs GP, Kreiger CJ, Luo D, Tian G, Garcia JS, Deleyrolle LP, et al. Glioma-derived CCL2 and CCL7 mediate migration of immune suppressive CCR2. Front Immunol. 2022;13:993444.

40. Li BH, Garstka MA, Li ZF. Chemokines and their receptors promoting the recruitment of myeloid-derived suppressor cells into the tumor. Mol Immunol. 2020;117:201–15.

41. Hein S, Müller V, Köhler N, Wikman H, Krenkel S, Streichert T, et al. Biologic role of activated leukocyte cell adhesion molecule overexpression in breast cancer cell lines and clinical tumor tissue. Breast Cancer Res Treat. 2011;129(2):347–60.

42. Kato Y, Tanaka Y, Hayashi M, Okawa K, Minato N. Involvement of CD166 in the activation of human gamma delta T cells by tumor cells sensitized with nonpeptide antigens. J Immunol. 2006;177(2):877–84.

43. Tan B, Shi X, Zhang J, Qin J, Zhang N, Ren H, et al. Inhibition of Rspo-Lgr4 Facilitates Checkpoint Blockade Therapy by Switching Macrophage Polarization. Cancer Res. 2018;78(17):4929–42.

44. Lu X, Li X, He Q, Gao J, Gao Y, Liu B, et al. miR-142-3p regulates the formation and differentiation of hematopoietic stem cells in vertebrates. Cell Res. 2013;23(12):1356–68.

45. Fordham JB, Naqvi AR, Nares S. Regulation of miR-24, miR-30b, and miR-142-3p during macrophage and dendritic cell differentiation potentiates innate immunity. J Leukoc Biol. 2015;98(2):195–207.

46. Wang WL, Ouyang C, Graham NM, Zhang Y, Cassady K, Reyes EY, et al. microRNA-142 guards against autoimmunity by controlling Treg cell homeostasis and function. PLoS Biol. 2022;20(2):e3001552.

47. Chiou GY, Chien CS, Wang ML, Chen MT, Yang YP, Yu YL, et al. Epigenetic regulation of the miR142-3p/interleukin-6 circuit in glioblastoma. Mol Cell. 2013;52(5):693–706.

48. Sun Y, Sun J, Tomomi T, Nieves E, Mathewson N, Tamaki H, et al. PU.1-dependent transcriptional regulation of miR-142 contributes to its hematopoietic cell-specific expression and modulation of IL-6. J Immunol. 2013;190(8):4005–13.

49. Wu LS, Lee SW, Huang KY, Lee TY, Hsu PW, Weng JT. Systematic expression profiling analysis identifies specific microRNA-gene interactions that may differentiate between active and latent tuberculosis infection. Biomed Res Int. 2014;2014:895179.

50. Xu G, Zhang Z, Wei J, Zhang Y, Guo L, Liu X. microR-142-3p down-regulates IRAK-1 in response to Mycobacterium bovis BCG infection in macrophages. Tuberculosis (Edinb). 2013;93(6):606–11.

