## Supplemental Table S1 for "Unraveling the Phenotype of Dormant Metastases Controlled by the Immune System"

**Supplementary Table S1.** Differentially expressed genes in Dormant metastases obtained by RNA-seq.

| Gene_id | Gene | Dormant-me | Nude-met | Over-met |
|---|---|---|---|---|
| | | log2*** | log2 | log2 |
| ENSMUSG000000288 | **Rspo1** | 10,0 | 0,0 | 0,4 |
| ENSMUSG000000337 | **Dysf** | 6,9 | 0,8 | 2,0 |
| ENSMUSG000000403 | **Arhgap9** | 6,4 | 0,4 | 2,2 |
| ENSMUSG000000258 | **Casp1** | 6,3 | 1,3 | 1,9 |
| ENSMUSG000000525 | **Dlg2** | 6,3 | 2,4 | -0,1 |
| ENSMUSG000000746 | **Foxs1** | 6,1 | 3,2 | 3,6 |
| ENSMUSG000000792 | **Slco1a6** | 6,1 | 2,2 | 0,7 |
| ENSMUSG000000744 | **S100a16** | 6,0 | 2,5 | 3,7 |
| ENSMUSG000000503 | **Ch25h** | 5,6 | -5,7 | 0,0 |
| ENSMUSG000000310 | **Smarca1** | 5,5 | 2,1 | -2,5 |
| ENSMUSG000000364 | **Lum** | 5,1 | 2,8 | -5,3 |
| ENSMUSG000000582 | **Kcnh1** | 4,6 | -3,1 | 1,2 |
| ENSMUSG000000064 | **Prl8a9** | 4,4 | -3,5 | -4,5 |
| ENSMUSG000000251 | **Bnc1** | 4,0 | -1,7 | -0,5 |
| ENSMUSG000000270 | **Lrp2** | 4,0 | 1,3 | 2,0 |
| ENSMUSG000000226 | **Alcam** | 3,9 | -3,4 | -4,2 |
| ENSMUSG000000707 | **Eid2b** | 3,8 | -2,9 | 0,2 |
| ENSMUSG000000283 | **Pappa** | 3,8 | -0,6 | -0,8 |
| ENSMUSG000000323 | **Pstpip1** | 3,6 | 1,5 | 0,3 |
| ENSMUSG000000283 | **Wdr31** | 3,6 | 0,4 | 1,9 |
| ENSMUSG000000177 | **Mmp9** | 3,5 | 0,8 | -1,3 |
| ENSMUSG000000435 | **Mdga1** | 3,4 | -1,6 | -4,5 |
| ENSMUSG000000989 | **Tmem185l** | 3,3 | -4,5 | -4,4 |
| ENSMUSG000000028 | **Cd97** | 3,1 | 0,4 | -2,5 |
| ENSMUSG000000390 | **Anpep** | 2,9 | 1,5 | -0,4 |
| ENSMUSG000000921 | **D030025P2** | 2,9 | -0,7 | 0,3 |
| ENSMUSG000000248 | **Gldc** | 2,9 | 1,0 | -1,4 |
| ENSMUSG000000994 | **RP23-480D** | 2,8 | -1,9 | -2,0 |
| ENSMUSG000000639 | **Slco1a5** | 2,7 | -0,4 | -0,3 |
| ENSMUSG000000316 | **Mfap3l** | 2,6 | 0,0 | -0,5 |
| ENSMUSG000000254 | **Adam8** | 2,5 | -1,1 | 1,0 |
| ENSMUSG000000248 | **Papss2** | 2,5 | -1,4 | -5,0 |
| ENSMUSG000000247 | **Fas** | 2,3 | -5,7 | 0,2 |
| ENSMUSG000000020 | **Ltbp2** | 2,2 | 0,1 | 0,6 |
| ENSMUSG000000461 | **Tmem229l** | 2,2 | -0,8 | -0,7 |
| ENSMUSG000000381 | **Slc43a2** | 2,2 | -0,4 | -0,5 |
| ENSMUSG000000557 | **Ghr** | 2,2 | -0,7 | -2,2 |
| ENSMUSG000000279 | **Vcam1** | 2,1 | 0,2 | 0,3 |
| ENSMUSG000000404 | **Blvrb** | 2,0 | -0,5 | -0,1 |

| | | | | |
|---|---|---|---|---|
| ENSMUSG000000174 | **Plxdc1** | 2,0 | -1,3 | -5,6 |
| ENSMUSG000000315 | **Slit2** | 2,0 | -0,9 | -2,8 |
| ENSMUSG000000353 | **Ccl7** | 2,0 | -2,9 | -2,5 |
| ENSMUSG000000414 | **Mfsd6** | 1,9 | -2,5 | -2,0 |
| ENSMUSG000000318 | **Cotl1** | 1,9 | -0,3 | -2,2 |
| ENSMUSG000000595 | **Calcrl** | 1,8 | -0,5 | -2,7 |
| ENSMUSG000000111 | **Adssl1** | 1,7 | 0,1 | -3,6 |
| ENSMUSG000000209 | **Ntn1** | 1,6 | 0,2 | 0,2 |
| ENSMUSG000000579 | **Gsta2** | 1,6 | -1,5 | -0,9 |
| ENSMUSG000000353 | **Ccl2** | 1,5 | -2,7 | -1,4 |
| ENSMUSG000000863 | **Gm12840** | 1,4 | -1,1 | -0,9 |
| ENSMUSG000000214 | **Serpinb9b** | 1,4 | -1,3 | -0,2 |
| ENSMUSG000000317 | **Mt2** | 1,4 | -1,1 | -3,0 |
| ENSMUSG000000186 | **Tbx3** | 1,4 | -1,2 | -2,9 |
| ENSMUSG000000323 | **Neo1** | 1,3 | -0,7 | -1,0 |
| ENSMUSG000000574 | **Phex** | 1,3 | -0,1 | -2,6 |
| ENSMUSG000000291 | **Pcdh7** | 1,3 | -0,7 | 0,0 |
| ENSMUSG000000478 | **A4galt** | 1,3 | -0,9 | -7,3 |
| ENSMUSG000000260 | **Stk17b** | 1,2 | -0,9 | 0,0 |
| ENSMUSG000000221 | **Dab2** | 1,2 | -0,3 | -3,7 |
| ENSMUSG000000216 | **Vcan** | 1,2 | -1,6 | -6,5 |
| ENSMUSG000000728 | **Ahnak2** | 1,2 | -0,5 | -0,6 |
| ENSMUSG000000320 | **Oaf** | 1,1 | -0,9 | -1,0 |
| ENSMUSG000000170 | **Slpi** | 1,1 | -2,2 | -3,0 |
| ENSMUSG000000376 | **Inf2** | 1,1 | -0,8 | -0,2 |
| ENSMUSG000000554 | **Srsf11** | -1,1 | 0,6 | 0,0 |
| ENSMUSG000000281 | **Zranb2** | -1,2 | 0,5 | 0,1 |
| ENSMUSG000000391 | **Whrn** | -1,2 | -2,7 | -3,8 |
| ENSMUSG000000246 | **Fads3** | -1,4 | 0,1 | 0,2 |
| ENSMUSG000000495 | **Cdc42ep1** | -1,9 | -0,3 | 0,6 |
| ENSMUSG000000240 | **Arhgap28** | -1,9 | 1,0 | 1,2 |
| ENSMUSG000000281 | **Adgrl2** | -2,0 | 0,5 | 1,0 |
| ENSMUSG000000288 | **Smpdl3b** | -2,3 | 0,6 | 0,5 |
| ENSMUSG000000416 | **Kif21b** | -2,4 | -0,3 | -0,2 |
| ENSMUSG000000245 | **Dpysl3** | -2,5 | -0,6 | 1,3 |
| ENSMUSG000000278 | **Shox2** | -2,6 | 0,0 | -0,3 |
| ENSMUSG000000548 | **Rnd1** | -2,8 | -0,5 | 0,7 |
| ENSMUSG000000266 | **Plxna2** | -3,1 | 0,2 | 1,1 |
| ENSMUSG000000033 | **Fkbp11** | -3,9 | -0,4 | 1,2 |
| ENSMUSG000000411 | **Hectd2** | -3,9 | 0,6 | 3,5 |
| ENSMUSG000000394 | **Prss23** | -5,4 | 0,9 | 0,4 |
| ENSMUSG000000080 | **Mid1ip1** | -5,7 | -0,3 | -1,4 |
| ENSMUSG000000226 | **Snai2** | -6,2 | 0,0 | -0,1 |

| | | | | |
|---|---|---|---|---|
| ENSMUSG000000790 | **Ly6c1** | -7,0 | -1,1 | 0,9 |
| ENSMUSG000000756 | **Ly6a** | -7,9 | -0,8 | 2,3 |
| ENSMUSG000000434 | **Otud1** | -8,5 | -0,9 | 0,8 |
| ENSMUSG000000315 | **Dlc1** | -8,8 | -0,9 | 2,0 |
| ENSMUSG000000318 | **Cdh13** | -9,2 | -0,1 | 1,6 |
| ENSMUSG000000708 | **Trabd2b** | -9,2 | -1,0 | 2,7 |
| ENSMUSG000000219 | **Ogdhl** | -9,5 | -0,4 | 0,2 |
| ENSMUSG000000281 | **Wls** | -9,9 | -0,8 | -1,3 |
| ENSMUSG000000470 | **Fbxo41** | -10,0 | 1,2 | 1,6 |
| ENSMUSG000000281 | **Depdc1a** | -10,0 | 1,0 | 0,0 |

*** $p < 0.001$ comparing Dormant-met group to the other two metastasis groups
